# Circuit and cellular mechanisms facilitate the transformation from dense to sparse coding in the insect olfactory system

**DOI:** 10.1101/240671

**Authors:** Rinaldo Betkiewicz, Benjamin Lindner, Martin P. Nawrot

## Abstract

Transformations between sensory representations are shaped by neural mechanisms at the cellular and the circuit level. In the insect olfactory system encoding of odor information undergoes a transition from a dense spatio-temporal population code in the antennal lobe to a sparse code in the mushroom body. However, the exact mechanisms shaping odor representations and their role in sensory processing are incompletely identified. Here, we investigate the transformation from dense to sparse odor representations in a spiking model of the insect olfactory system, focusing on two ubiquitous neural mechanisms: spike-frequency adaptation at the cellular level and lateral inhibition at the circuit level. We find that cellular adaptation is essential for sparse representations in time (temporal sparseness), while lateral inhibition regulates sparseness in the neuronal space (population sparseness). The interplay of both mechanisms shapes dynamical odor representations, which are optimized for discrimination of odors during stimulus onset and offset. In addition, we find that odor identity is stored on a prolonged time scale in the adaptation levels but not in the spiking activity of the principal cells of the mushroom body, providing a testable hypothesis for the location of the so-called odor trace.

## 1 Introduction

How nervous systems process sensory information is a key issue in systems neuroscience. Animals are required to rapidly identify behaviorally relevant stimulus features in a rich and dynamic sensory environment, and neural computation in sensory pathways is tailored to this need. Sparse stimulus encoding has been identified as an essential feature of sensory processing in higher brain areas in both, invertebrate [1, 2, 3, 4, 5] and vertebrate [6, 7, 8, 9] systems. Sparse representations provide an economical means of neural information coding [10, 11] where information is represented by only a small fraction of all neurons (population sparseness) and each activated neuron generates only few action potentials (temporal sparseness) for a highly specific stimulus configuration (lifetime sparseness).

The nervous systems of insects have limited neuronal resources and thus require particularly efficient coding strategies. The insect olfactory system is analogue to the vertebrate olfactory system and has become a popular model system for the emergence of a sparse code. We use a computational approach to study the transformation from a dense olfactory code in the sensory periphery to a sparse code in the mushroom body (MB), a central structure of the insect brain important for multimodal sensory integration and memory formation. A number of recent studies emphasized the role of sparse coding in the MB. In locusts, sparse responses were shown to convey temporal stimulus information [12]. In Drosophila, sparse coding was found to reduce overlap between odor representations and facilitate their discrimination [13]. Consequently, sparse coding is an essential feature of plasticity models for olfactory learning in insects [14, 15, 16, 17, 18] and theoretical work has emphasized the analogy of the transformation from a dense code in projection neurons (PNs) to a sparse code in Kenyon cells (KCs) with dimensionality expansion in machine learning methods [14, 19, 20].

Central to our modeling approach are two fundamental mechanisms of neural computation that are ubiquitous in the nervous systems of invertebrates and vertebrates. Spike-frequency adaptation (SFA) is a cellular mechanism that has been suggested to support efficient and sparse coding and to reduce variability of sensory representation [21, 22, 23]. Lateral inhibition is a basic circuit design principle that exists in different sensory systems, mediates contrast enhancement and facilitates stimulus discrimination [24, 25, 26, 27]. Both mechanisms are evident in the insect olfactory system. Responses of olfactory receptor neurons (ORNs), local interneurons (LNs) and PNs in the antennal lobe (AL) show stimulus adaptation [28, 29] and strong adaptation currents have been identified in KCs [30, 31]. Lateral inhibition in the AL is mediated by inhibitory LNs [32]. It is crucial for establishing the population code at the level of PNs [29, 33], for gain control [34, 35], for decorrelation of odor representations [36], and for mixture interactions [29, 37, 38].

Taken together, we find that lateral inhibition and spike-frequency adaptation account for the transformation from a dense to sparse coding, decorrelate odor representations, and facilitate precise temporal responses on short and long time scales.

## 2 Results

### Spiking network model of the olfactory pathway with lateral inhibition and spike-frequency adaptation

We designed a spiking network model that reduces the complexity of the insect olfactory processing pathway to a simplified three-layer network (Fig. 1A) that expresses the structural commonality across different insect species: an input layer of olfactory receptor neurons (ORNs), subdivided into different receptor types, the AL, a first order olfactory processing center, and the MB. Furthermore, the model combines two essential computational elements: (i) lateral inhibition in the AL, and (ii) spike-frequency adaptation in the AL and the MB.

**Fig. 1.**
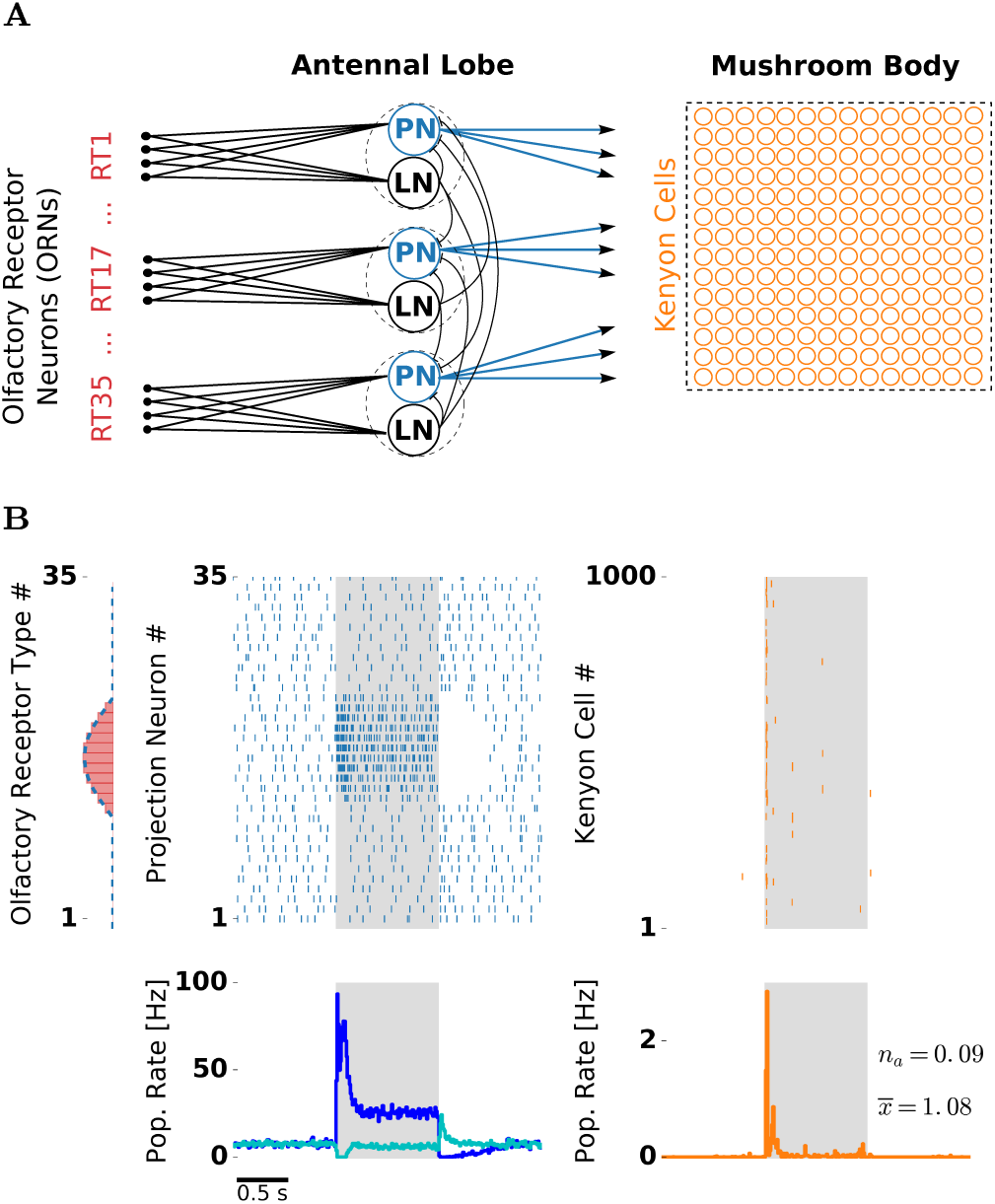
Olfactory network model structure and odor response. *(A)* Network structure resembles the insect olfactory pathway with three main processing stages. In each glomerulus (dashed circles), a PN (blue) and a LN receive convergent ORN input (red) by one receptor type (RT). Each LN provides unspecific lateral inhibition to all PNs. KCs (orange) receive on average 12 inputs from randomly chosen PNs. *(B)* Receptor response profile (red bars; AL input) depicts the evoked firing rate for each ORN type. Evoked PN spike counts (dashed blue line; AL output) follow the ORN activation pattern. Raster plots depict single trial responses of PNs (blue) and KCs (orange). Presentation of an odor during 1000 ms is indicated by the shaded area. Population firing rates were obtained by averaging over 50 trials. PN spikes display a temporal structure that includes evoked transient responses at stimulus on- and offset, and a pronounced inhibitory post-odor response. PN population rate was averaged over PNs showing “on” responses (blue) and “off” responses (cyan). KC spikes were temporally sparse with majority of the spikes occurring at the stimulus onset.

The processing between the layers is based on excitatory feedforward connections. Converging receptor input from all ORNs of one type is received by spatially confined subunits of the AL called glomeruli. In our model, glomeruli are represented by a single uniglomerular PN and a single inhibitory local interneuron (LN). In the MB, each KC receives on average 12 PN inputs [2], based on a random connectivity between the AL and the MB [39]. All neurons in the AL and the MB were modeled as leaky integrate-and-fire neurons with spike-triggered adaptation. Based on evidence from theoretical [40] and experimental studies [41], adaptation channels cause slow fluctuations. We accounted for this fact by simulating channel noise in the slow adaptation currents (cf. Methods).

We simulated ORN responses to different odor stimuli. Single ORN responses were modeled in the form of Poisson spike trains with firing rates dependent on the receptor type and stimulus identity. The relationship is set by a receptor response profile (Fig. 1B left) which determines ORN firing rates of all receptor types for a given stimulus. Responses to different stimuli are generated by shifting the response profile along the receptor space (Fig. 2). The offset between any two stimuli reflects their dissimilarity – similar stimuli activate overlapping sets of olfactory receptors, whereas dissimilar stimuli activate largely disjoint sets of receptors. Stimuli were presented for one second, reflected by a step-like increase of ORN firing rate.

**Fig. 2.**
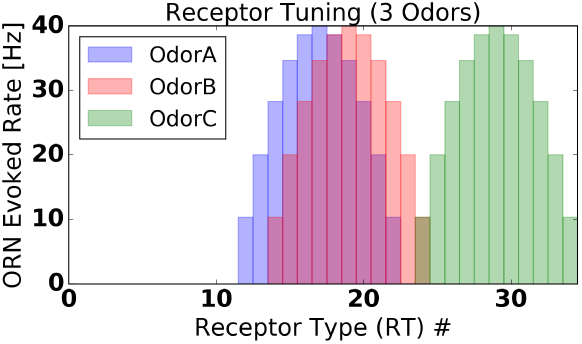
Receptor response profile for two similar odors (red, blue) and a dissimilar odor (green).

In the absence of stimuli, ORNs fired with a rate of 20 Hz reflecting their spontaneous activation [28]. Both LNs and PNs receive direct ORN input. We tuned synaptic weights of the model to match physiologically observed firing rates of PNs and LNs, which are both about 8 Hz [1, 42, 43] (for details see Methods). Lateral inhibition and spike-frequency adaptation, the neural mechanisms under investigation, both provide an inhibitory contribution to a neuron’s total input. In our model, spike-frequency adaptation is a cellular mechanism mediated by a slow, spike-triggered, hyperpolarizing current in LNs, PNs and KCs, whereas a global lateral inhibition in the AL is mediated by LNs with fast synapses that receive input from a single ORN type and inhibit all PNs in a uniform fashion.

### Odor responses at the AL and the MB level of the spiking network model

Figure IB illustrates PN and KC responses to one odor. PNs driven by the stimulus showed a strong transient response at the stimulus onset, a pronounced adaptation during the stimulus, and a period of silence after stimulus offset due to the slow decay of the strong adaptation current. This resembles the typical phasic-tonic response patterns of PNs [43, 44].

PNs receiving direct input from ORNs activated by the stimulus, showed a strong response at the stimulus onset. Interestingly, the population firing rate over these PNs revealed that the “on” response follows a biphasic profile with an early and a late component. In addition, PNs with no direct input from stimulated ORNs showed an “off” response at the stimulus offset. Non-driven PNs were suppressed during a short period after stimulus onset, and showed reduced firing during the tonic response. The PN population response consisted of complex activations of individual PNs with phases of excitation and inhibition. Hence, in the AL, odors were represented as spatio-temporal spike patterns across the PN population.

At the level of the MB, KCs typically show none or very little spiking during spontaneous activity and respond to odors with only a few spikes in a temporally sparse manner [1, 3, 4]. In our model, synaptic weights between PNs and KCs were tuned to match the very low probability of spontaneous firing. Resulting KC responses were temporally sparse. Due to the negative feedback mediated by strong spike-frequency adaptation, most KC spikes were confined to stimulus onset. Notably, we also found that KCs sometimes exhibited “off” responses. These KC “off” spikes occurred very rarely, because they are driven by the PN “off” response, which is much weaker compared to the PN “on” response. Timing and amplitude of temporally sparse responses are in good quantitative agreement with in vivo KC recordings [3].

### Dense and dynamic odor representations in the AL

In order to explore effects of lateral inhibition and cellular adaptation on stimulus representations, we simulated odor responses in conditions in which we separately deactivated one or both mechanisms. Lateral inhibition was deactivated by setting the inhibitory synaptic weight between LNs and PNs to zero and simultaneously reducing the value of the excitatory synaptic weight between ORNs and PNs, such that the spontaneous firing rate of 8 Hz was kept. Adaptation was deactivated by replacing the dynamic adaptation current by a constant current with an amplitude that maintained the average spontaneous firing rate.

Figure 3 illustrates the separate effects of lateral inhibition and adaptation on odor responses in the PN population. In all conditions, PNs fired spontaneously before stimulation due to spontaneous ORN activation. PNs driven by stimulation receive input from ORNs that were activated by the presented odor. In the absence of adaptation and lateral inhibition (Fig. 3 (i)) the stimulus response followed the step-like stimulation and showed no further temporal structure. In the presence of lateral inhibition (Fig. 3 (ii)), PNs not driven by the stimulus were strongly suppressed. Adaptation alone (Fig. 3 (iii)) resulted in a phasic-tonic response profile with a high phasic peak amplitude immediately after stimulus onset. In the presence of both mechanisms (Fig. 3 (iv)) we observed the characteristic phasic-tonic PN response. The transient response was reduced in peak amplitude, and, interestingly, followed a biphasic profile with an early and a late component.

**Fig. 3.**
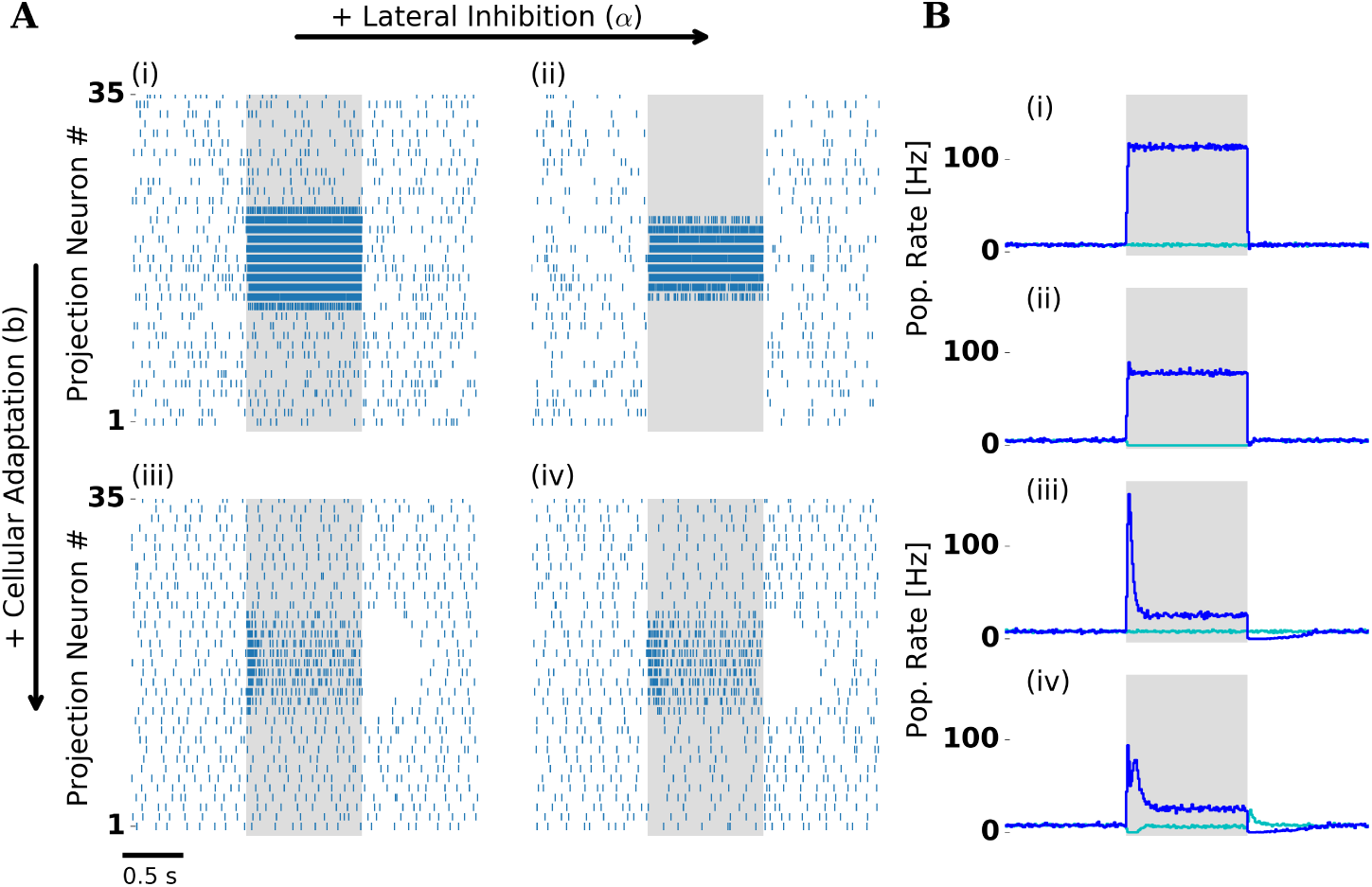
Lateral inhibition and cellular adaptation shape PN odor response dynamics. *(A)* Single trial PN spiking responses simulated with (right column) and without (left column) lateral inhibition, and with (bottom row) and without (top row) adaptation. Presentation of a single odor during 1000 ms is indicated by the shaded area. With adaptation PNs display a temporal structure that includes a transient and a tonic response, and a pronounced inhibitory post-odor response. *(B)* Trial averaged population firing rate: PNs driven by stimulation (blue) and remaining PNs (cyan). Panels (i)-(iv) indicate presence and absence of lateral inhibition and adaptation as in (A). In the presence of lateral inhibition firing rates during stimulation are reduced. In the presence of lateral inhibition and adaptation (iv) PNs show either transient “on” responses (blue) or “off” responses (cyan). Panels A (iv) and B (iv) are reproduced in Fig. IB.

In our model, the interaction of lateral inhibition and the intrinsic adaptation currents in LNs and PNs accounts for biphasic PN responses. Because lateral inhibition is strongest at stimulus onset, most of the phasic PN response was delayed (late component), whereas the immediate PN response (early component) was not affected. Fast and delayed PN responses have also been found experimentally in the honeybee [45].Model evidence for the interplay of cellular and network mechanisms behind biphasic PN responses was found in the pheromone system of the moth [46].

### Spike-frequency adaptation supports temporal sparseness in the MB

To isolate the contributions of adaptation and lateral inhibition (the latter present only at the AL level) to odor responses at the MB level, we again tested the four conditions by deactivating one or both mechanisms. In all four conditions, KCs were almost silent and spiked only sporadically during spontaneous activity, whereas amplitude and temporal profile of their odor response differed across conditions (Fig. 4).

**Fig. 4.**
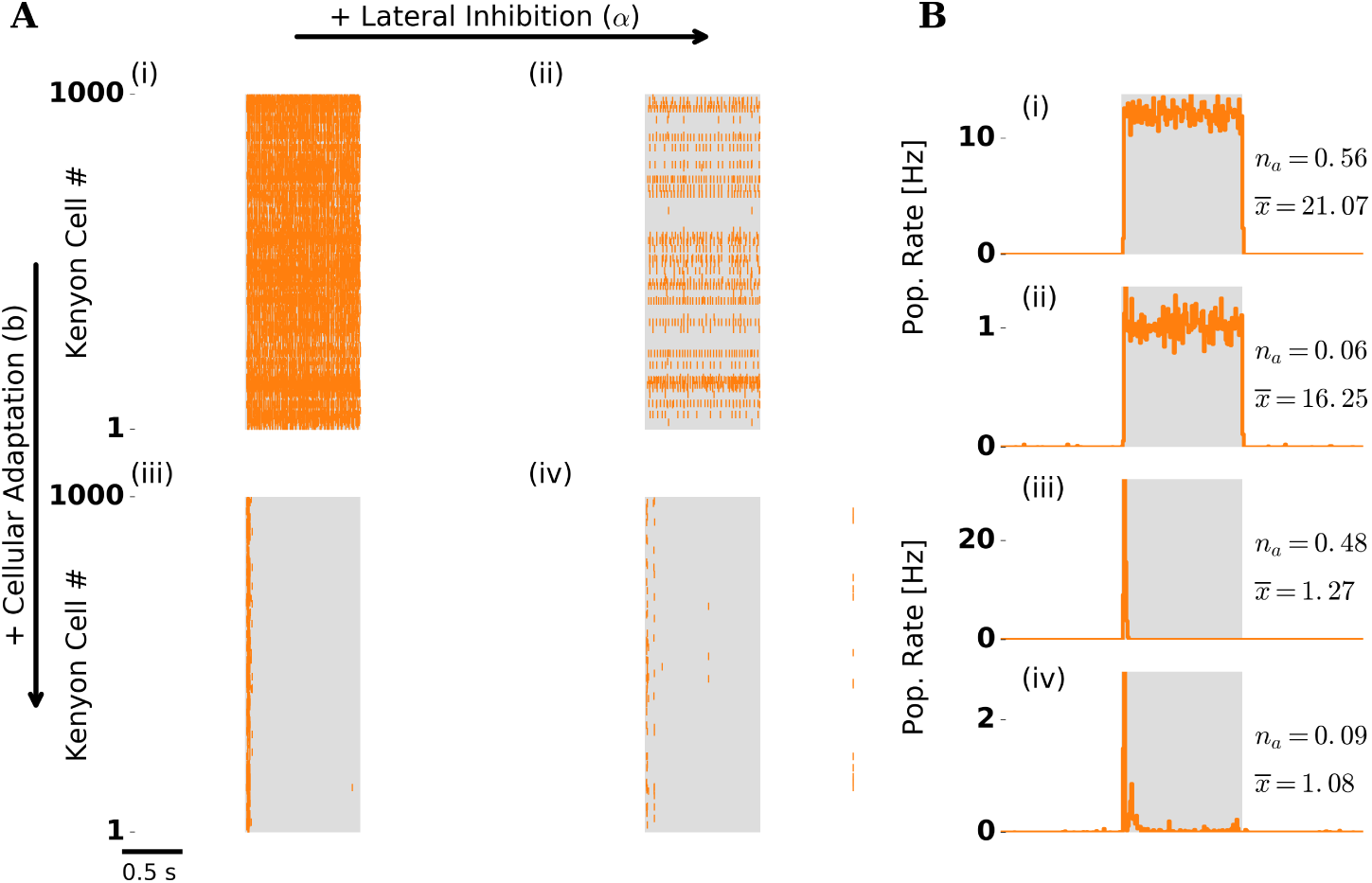
Odor response dynamics of the KC population. Figure layout as in Fig. 3. *(A)* Single trial population spike raster responses. *(B)* Trial averaged KC population firing rate. Numbers to the right indicate the fraction of activated KCs *(n_a_)* and the mean number of spikes per activated KC during stimulation 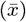. Without adaptation (i,ii) KCs spike throughout stimulation because PN drive is strong and persistent. The fraction of activated KCs drops in the presence of lateral inhibition (ii,iv). With adaptation (iii,iv) most of KC spikes are confined to the stimulus onset, indicating temporally sparse responses. Panels A (iv) and B (iv) are reproduced in Fig. 1B.

In the presence of adaptation we observed temporally sparse responses (Fig. 4 (iii)–(iv)). KCs typically responded with only 1–3 spikes (mean spikes per responding KC were slightly above one, compare 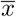 in Fig. 4B (iii),(iv)). Due to the negative feedback mediated by strong spike-frequency adaptation, most KC spikes were confined to stimulus onset.

In the absence of adaptation and regardless of the presence (Fig. 4 (i)) or absence (Fig. 4 (ii)) of lateral inhibition, responding KCs fired throughout stimulation, because they received persistently strong input from PNs. Such persistent KC responses are in disagreement with experimental observations [1, 3, 4].

We quantified temporal sparseness of KC responses by calculating a measure modified from [47] (cf. Methods). Comparison of temporal sparseness across the four conditions confirms that KC responses were temporally sparse only in the presence of adaptation whereas lateral inhibition had no effect on temporal sparseness (Fig. 5A).

**Fig. 5.**
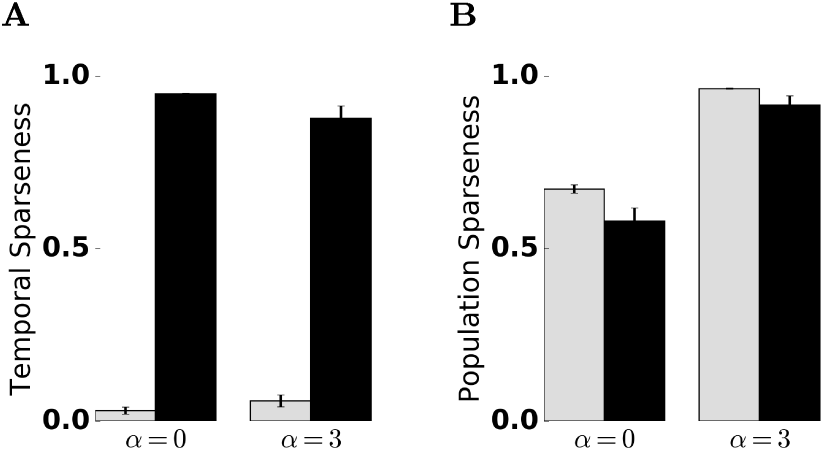
Quantification of temporal and population sparseness in the KC population. Sparseness was measured in the absence (*α* = 0) and presence (*α* = 3) of lateral inhibition, and the presence (black bars) and absence (gray bars) of spike-frequency adaptation. The sparseness measure was averaged over 50 trials. Error bars indicate standard deviation. A value of one corresponds to maximally sparse responses. *(A)* Adaptation promotes temporal sparseness. *(B)* Lateral inhibition in the AL promotes KC population sparseness.

### Lateral inhibition supports population sparseness in the MB

We observed that the fraction of responding KCs was considerably lower in the presence of lateral inhibition (compare *n_a_* across panels in Fig. 4B). We recall that lateral inhibition in our model is acting on PNs. A reduced PN population rate caused by lateral inhibition (compare Fig. 3 (ii),(iv)) is reflected in a lower net input to KCs. How does this affect KC responses on a population level?

We visualized MB odor representations with activation patterns obtained by arranging KC spike counts evoked by two similar odors on a 30×30 grid in arbitrary order (Fig. 6A). In the absence of lateral inhibition (Fig. 6A top), a majority of the KC population was activated by both tested odors, due to strong PN input. KCs responded with 1–3 spikes. In the presence of lateral inhibition (Fig. 6A bottom), the fraction of activated KCs was reduced substantially (KCs activated, trial averaged: 9%, std: 3%), whereas the range of individual KC responses (1–3 spikes) was not affected. These activation patterns demonstrate that the MB odor representations are sparse on a population level, as each odor is represented by the activity of a small fraction of the KC population.

**Fig. 6.**
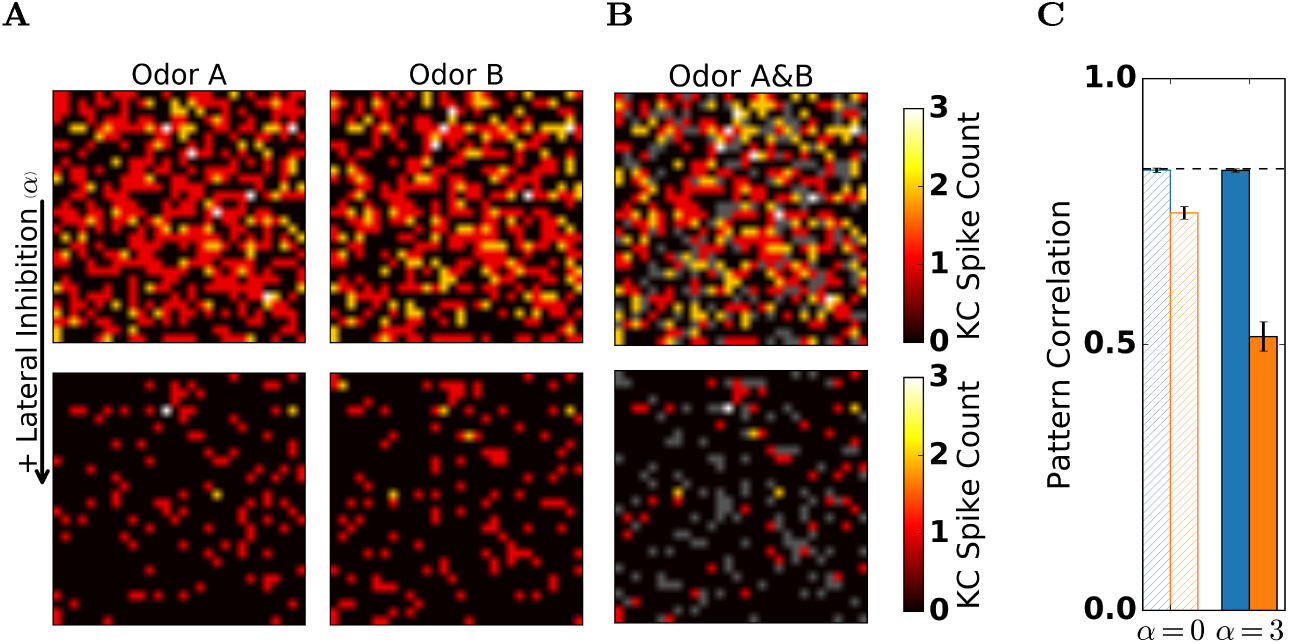
Lateral inhibition in the AL facilitates population sparseness and reduces pattern correlation in the MB. Spike counts (single trial) of 900 randomly selected KCs in response to two similar odors (“Odor A” and “Odor B”) arranged on a 30×30 grid in the absence (top row) and in the presence (bottom row) of lateral inhibition. Inactive KCs are shown in black. *(A)* In the absence of lateral inhibition KCs readily responded to both odors, resulting in an activation pattern where most KCs are active. In the presence of lateral inhibition both odors evoked sparse KC activation patterns. *(B)* Superposition of responses to the two odors. KCs that were activated by both odors are indicated by hot colors (color bar denotes spike count of the stronger response). KCs that were activated exclusively by one of the two odors are indicated in gray. The fraction of KCs that show overlapping responses is reduced in the presence of lateral inhibition. (C) Pattern correlation between responses to the two odors obtained for PN (blue) and KC (orange) spikes counts, in the absence (*α* = 0) and presence (*α* = 3) and of lateral inhibition. Dashed line indicates pattern correlation of the input (ORNs). Pattern correlation was retained at the AL and reduced at the MB level. Lateral inhibition in the AL reduced pattern correlation in KCs but not in PNs.

To quantify population sparseness of odor representations in the MB, we again calculated a sparseness measure (cf. Methods). Population sparseness increased in the presence of lateral inhibition, independent of spike-frequency adaptation (Fig. 5B). In the presence of lateral inhibition and spike-frequency adaptation, both population and temporal sparseness were in qualitative and quantitative agreement with experimental findings [1, 2, 3, 4]. Taken together, odor representations at the MB level were characterized by a small fraction of the KC population responding with a small number of spikes.

### Decorrelation of odor representations between AL and MB

In our model, lateral inhibition in the AL increased population sparseness of MB odor representations. Given sparse population responses, does the overlap between MB odor representations decrease? We visualized the overlap between odor representations in the MB by overlaying KC activation patterns in response to two similar odors (Fig. 6B). With lateral inhibition, most of the KC responses were unique to odor A or odor B (shown in gray in Fig. 6B) and only relatively few KCs were activated by both odors. In contrast, with lateral inhibition deactivated (Fig. 6B top), the ratio of KCs with unique responses (gray) to the total number of activated cells (all colors) was low, indicating highly overlapping responses.

We quantified the overlap between odor representations evoked by two similar odors in the PN and the KC population. To this end, we calculated Pearson’s correlation coefficient between spike counts evoked by either odor, across the corresponding population (Fig. 6C, cf. Methods). Interestingly, PNs retained the overlap of the input, independent of lateral inhibition. In contrast, KC representations showed a reduced overlap that decreased even further in the presence of lateral inhibition.

We tested how scaling of the lateral inhibition strength affected the pattern overlap in PN and KC odor representations. To this end, we varied the strength of lateral inhibition (*α*) in the AL by increasing the strength of inhibitory synapses and adjusting feedforward weights (see Methods). In addition, we calculated pattern correlations in the absence of adaptation. As before, pattern correlation was calculated for two similar odors that activated an overlapping set of receptors (cf. Fig. 2). In the absence of adaptation, lateral inhibition robustly decorrelated odor representations in both populations (Fig. 7B). In the presence of adaptation, increasing lateral inhibition had different effects on the PN and KC population (Fig. 7A). In PNs the correlation of the input was retained for all tested values of lateral inhibition. In KCs pattern correlation first decreased for weak to moderate lateral inhibition strength but then increased for strong lateral inhibition. For an intermediate strength of the inhibitory weights the pattern correlation between KC responses to similar odors attained a minimal value. In general, a reduction of pattern correlation from PN to KC representations is characteristic for the insect MB [48]. Furthermore low overlap between KC representations has been found to facilitate discrimination of odors [49]. We therefore choose the intermediate strength of the inhibitory weights (α = 3) as a reference point in our model.

**Fig. 7.**
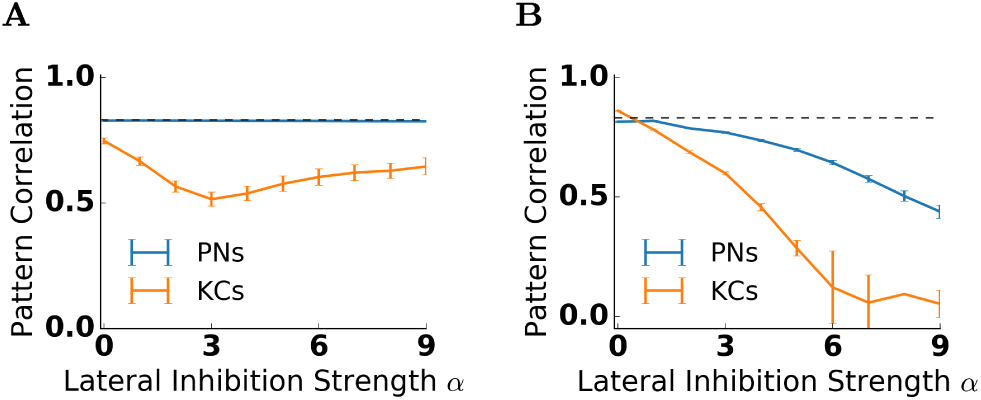
Pattern correlation in the antennal lobe and the mushroom body depend on lateral inhibition strength *α*. The correlation coefficient between the response patterns to two similar odors was calculated and averaged over 50 trials for PNs *(blue)* and KCs *(orange).* Error bars indicate standard deviation. Pattern correlation of the input is indicated by the dashed line. Input correlation is high because similar odors activate largely overlapping set of receptors. *(A)* In the presence of adaptation, pattern correlation in PNs *(blue)* stays close to the input correlation for all values of lateral inhibition strength. In KCs *(orange)* the correlation decreases for small values of lateral inhibition strength, and increases for large values of lateral inhibition strength. Pattern correlation in KCs is minimal for *α* = 3. *(B)* In the absence of adaptation, pattern correlation decreases with the lateral inhibition strength both in PNs and KCs. The decrease is stronger in KCs.

### Odor encoding on short and long time scales

Next, we tested if in our model the information about stimulus identity is contained in AL and MB odor representations, by performing a decoding analysis in subsequent time bins of 50 ms (cf. Methods). In PNs decoding accuracy peaked during stimulus on- and offset (Fig. 8A). Both peaks coincide with a state of transient network activity caused by the odor on-or offset. The “on” and the “off” responsive PNs establish odor representations optimized for discrimination. After stimulus onset, decoding accuracy dropped but remained on a plateau well above chance level. Remarkably, after stimulus offset, odor identity could be decoded for an extended time period (several hundreds of ms) albeit with a reduced accuracy. Such odor after effects have been demonstrated previously in experiments [50, 44](cf. Discussion).

**Fig. 8.**
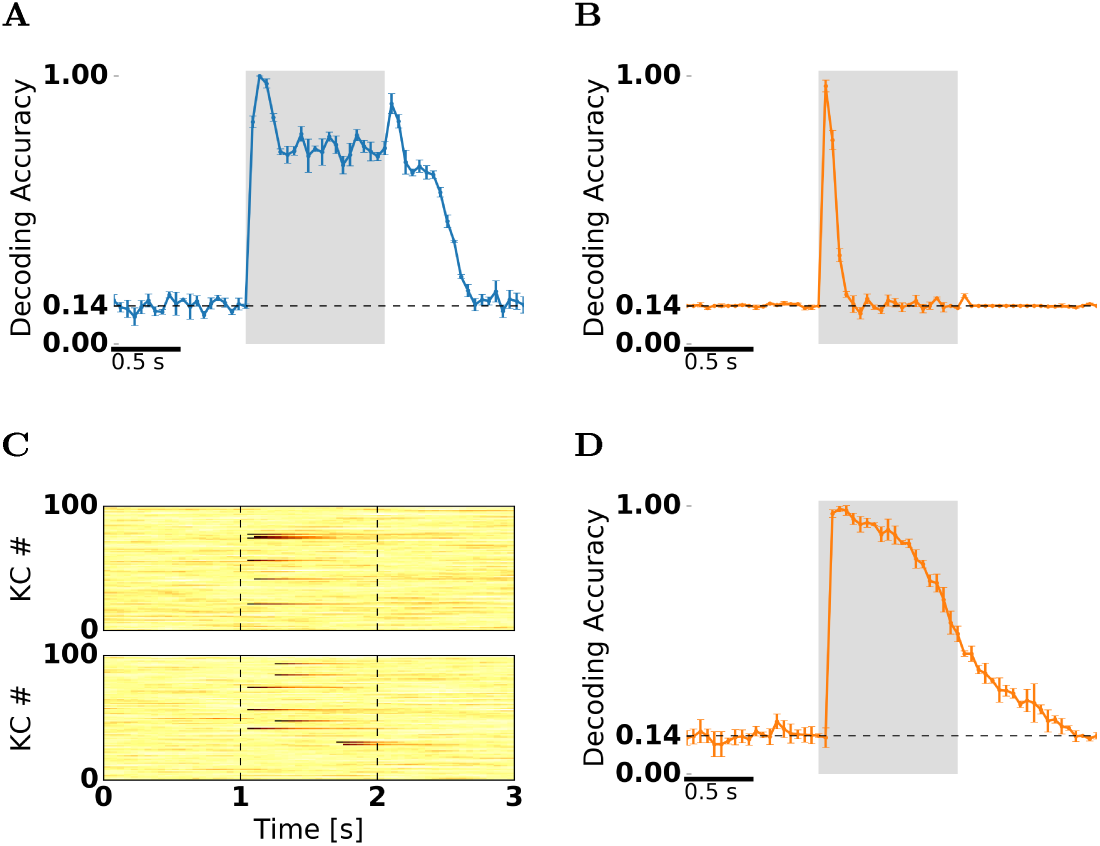
Decoding of odor identity indicates a prolonged and reliable odor information in KC adaptation currents. *(A,B,D)* Decoding accuracy was calculated for non-overlapping 50 ms time bins, based on a set of seven stimuli (chance level ≈ 0.14) presented for one second (shaded area). Blue shading indicates standard deviation obtained from a cross-validation procedure (see Methods). *(A)* Decoding of odor identity from PN spike counts. Decoding accuracy peaks at odor on- and offset, and remains high after stimulation. *(B)* Decoding of odor identity from KC spike counts. Decoding accuracy is above chance only in the first three bins following stimulus onset. *(C)* Adaptation current amplitudes (single trial, hot colors in arbitrary units) of 100 selected KCs in response to “odor A” (top) and “odor B” (bottom). *(D)* Decoding of odor identity from KC adaptation currents. Decoding accuracy peaks 150 ms after odor onset, then drops during stimulation but remains high and is sustained after odor offset.

In KCs decoding accuracy was above chance level only in the first 2–3 time bins (about 100 ms) after stimulus onset (Fig. 8B). In all other time bins decoding accuracy remained at chance level. Because the spiking activity in the KC population is temporally sparse, the continuous information at the AL output is lost in the MB spike count representation. This raises the question whether and if so how the information could be preserved in the MB throughout the stimulus. The intrinsic time scale of the adaptation currents might potentially support prolonged odor representations (Fig. 8C). We therefore repeated the decoding analysis on the adaptation currents measured in KCs (Fig. 8D). Indeed, the stimulus identity could reliably be decoded based on the intensity of the adaptation currents in subsequent time bins of 50 ms. Decoding accuracy peaked after stimulus onset and then slowly decreased. Remarkably, because KCs show very little spontaneous activity, the decay of the classification performance in the absence of stimulation, is caused by slow adaptation current fluctuations due to channel noise.

## 3 Discussion

We investigated the transformation between dense AL and sparse MB odor representations in a spiking network model of the insect olfactory system. Our generic model demonstrates lateral inhibition and spike-frequency adaptation as sufficient mechanisms underlying dynamic and combinatorial responses in the AL that are transformed into sparse MB representations. To simulate responses to different odors we incorporated simple ORN tuning and glomerular structure in our model. This approach allows us to investigate how different odors are represented in the AL and MB population activity and asses information about odor identity contained in respective odor representations. We inspected overlap between odor representations in both populations. Sparse coding reduces overlap between representation, as has been predicted on theoretical grounds [51, 52, 53] and shown for MB odor representations [2, 4, 13]. Similarly, our model shows pattern decorrelation in the MB but not in the AL.

### Post-odor responses

In our model, we found “on” and “off” responsive PNs. At the stimulus offset, the “off” responsive PNs transiently increase, whereas the “on” responsive PNs transiently decrease their firing rate (cf. Fig. 3). “On” responsive PNs remain adapted beyond stimulus offset.Their excitability thus stays reduced until the slow adaptation current has decayed. In contrast, in “off” responsive PNs increased lateral inhibition during stimulation causes a below-baseline adaptation level throughout the stimulus and thus an increased excitability. In effect, the odor-evoked and the post-odor PN activation patterns are reversed, i.e. anti-correlated (not shown). This result matches well the experimental observations in honeybee [50, 54, 44] and *Drosophila* [55] PNs. Those results show highly correlated response patterns throughout stimulation, and stable but anti-correlated post-odor response patterns.

### Differential mechanism underlying temporal and population sparseness in KCs

In our model, the two mechanisms underlying temporal sparseness and population sparseness are independent.

Temporal sparseness of KC responses in our model compares well to the experimentally recorded spiking responses in *Drosophila*, locust and moth [1, 3, 4], and to calcium imaging experiments in the honeybee [2]. The model proposed here solely relies on spike-frequency adaptation for temporally sparse responses. On a cellular level, strong adaptation currents in KCs, which are suitable for generation of sparse responses, have been found in the honeybee [30] and cockroach [31]. In our model temporal sparseness is not affected by the deactivation of lateral inhibition, a finding supported a previous study [23].

Several studies have suggested that either feedforward inhibition [56] or feedback inhibition [2, 57, 12, 58, 59] causes temporally sparse responses. The existence of inhibitory feedback neurons in the MB has been demonstrated experimentally in different insect species (cockroach [60], *Drosophila* [61], honeybee [62], locust [57]), whereas evidence for feedforward inhibition to the MB is lacking [12]. Our model demonstrates that temporally sparse responses can be obtained without an inhibitory circuit motive. There is further evidence for a GABA-independent mechanism for the temporal shortening of KC responses. Calcium imaging studies in *Drosophila* [58, 13] and in the honeybee [63] showed that the temporal profile of KCs’ fast response dynamics is preserved independent of GABA inhibition. In addition, cellular mechanism such as high threshold for KC activation in *Drosophila* [4] and active KC subthreshold properties in locust [1, 64] have been shown to support population sparseness.

The KC population sparseness in our model matches qualitatively and quantitatively with experimental estimates from electrophysiological responses in locust and *Drosophila* [1, 4] and from calcium imaging in *Drosophila* [5]. Our model shows sparse KC responses on a population level in the presence but not in the absence of lateral inhibition. Calcium imaging experiments in the honeybee [63, 23] have shown that inactivating GABA transmission disrupts population sparseness, in line with our modeling results. In *Drosophila* feedback inhibition contributes to the population sparseness of KCs, as blocking of feedback inhibition reduced population sparseness and undermined the learned discrimination of similar odors [58, 13]. In addition, plasticity of inhibitory feedback changing response patterns in the KC population might be crucial for associative learning [61, 65, 66, 67]. We suggest that different mechanism of sparseness are not mutually exclusive. Both lateral inhibition in the AL and feedback inhibition in the MB are likely to be necessary for sparse KC population responses.

### Decorrelation of odor representations between AL and MB

Decorrelation of stimulus representations has been postulated to be a fundamental principle underlying sensory processing [68, 69]. In particular, in the olfactory system odor representations are transformed to decorrelate activity patterns evoked by similar odors making them more distinct [70, 71, 72]. Transformations decreasing the overlap between representations are termed pattern decorrelation. Less overlapping representations increase memory capacity [47] and make discrimination of odors easier [49]. In our model, we found that AL odor representations preserved the similarity of the input, whereas representations of similar odors at the periphery became decorrelated in the MB.

We quantified the effects of lateral inhibition and adaptation on pattern correlations. We found that, in the AL decorrelation of activity patterns occurred only in the absence of adaptation. Moreover, the amount of decorrelation depended on lateral inhibition strength. In computational studies lateral inhibition was previously shown to decorrelate odor representations [73, 19]. In a *Drosophila* study using extracellular recording, lateral connection in the AL were found not to affect correlations between glomerular channels [74], but there is also evidence for decorrelation of AL representations [75]. In our model, pattern correlation between representations of similar odors was preserved at the level of the AL but reduced in the MB.

### Odor representation in adaptation currents

Early investigations of dynamical odor representations have shown that odor identity can be reliably decoded from PN spike counts in 50 ms time bins [34, 76]. We used this approach to show that odor representations were specific and reliable in our model, including both AL and MB odor representations. We found that at the AL level, odor representation were optimized for discrimination during odor on- and offset. In line with previous findings in PNs [76, 29] the peak accuracy coincided with transient network activity. Unlike in the AL, at the MB level, stimulus identity could be decoded from KC spike counts only during a short time window after stimulus onset (up to about 150 ms, see Fig. 8B). This is a consequence of the temporally sparse KC responses.

Moreover, we found that KC adaptation currents retain a representation of stimulus identity, resembling a prolonged odor trace [77, 78]. In our model, an odor trace present in adaptation levels extends well beyond the brief spiking responses. Adaptation currents constitute an internal dynamical state of the olfactory network that is not directly accessible to downstream neurons – a “hidden state” [79]. However, adaptation levels influence the responses to (subsequent) stimuli [23] and may also affect downstream processing through an indirect pathway.

Our results suggest that odor representations are not exclusively found in the spiking activity. Calcium and calcium-dependent adaptation currents were found in *cockroach* and honeybee KCs [30, 31]. Therefore, an odor trace in KC adaptation levels should be reflected in their intracellular calcium concentration. Odor representations in the calcium signal are likely to mediate the formation of associative memories through biochemical mechanisms on the cellular level. We predict long-lasting levels of calcium in the KC population that retain odor information and provide a potential substrate for a short-term sensory memory. Therefore, classification of calcium levels recorded in the MB should reveal odor identity on a time scale determined by the decay of the intracellular calcium level. Indeed, a recent study by [80] showed that prolonged calcium activity in KCs encoded odor information and could be related to behavioral odor recognition performance in trace conditioning experiments.

## 4 Methods

### 4.1 Spiking network model

A spiking network model with 3 layers (ORN, AL and MB, see Fig. 1AB) was simulated using Brian 1.4 [81]. The model includes 35 ORN types, 284 ORNs per type, 35 PNs and LNs, and 1000 KCs. Each of the 35 LN-PN pairs constitute a glomerulus. Across insect species, the number of glomeruli varies from a few tens to several hundred, we based our model on the lower end of this range. The ratio between the number of PNs and KCs is roughly based on the data available in *Drosophila* [4].

The connections between the 3 network layers (ORNs, AL, MB) are feed-forward and excitatory. Within the AL, LNs provide lateral inhibition to PNs. ORNs provide input to PNs and LNs. All ORNs of the same receptor type target the same, single glomerulus. Every LN has inhibitory connections with all PNs, mediating unspecific lateral inhibition within the AL. Every KC receives 12 PN inputs on average [2]. Connections between PNs and KCs were randomly drawn. Synaptic weights between all neurons are given in Table 1 for four different simulation conditions (cf. Sec. 4.1.3).

**Tab. 1.** Synaptic weights for *w_OL_* (ORN-LN), *w_Op_* (ORN-PN), *w_LP_* (LN-PN) and *w_PK_* (PN-KC) connections in different simulation conditions ((i)-(iv)).

|  | (i) | (ii) | (iii) | (iv) |
| --- | --- | --- | --- | --- |
| $w_{OL}$ | 1 nS | 1 nS | 1 nS | 1 nS |
| $w_{OP}$ | 1 nS | 1.12 nS | 1 nS | 1.12 nS |
| $w_{LP}$ | 0 nS | 3 nS | 0 nS | 3 nS |
| $w_{PK}$ | 5 nS | 5 nS | 5 nS | 5 nS |

Responses to a set of 7 stimuli, 50 trials each, and 3000 ms trial duration were simulated. Stimuli had a duration of 1000 ms and were presented at t=1000 ms. To ensure steady state initial conditions, simulations were initialized for 2000 ms without recording the activity.

#### 4.1.1 Receptor input

ORNs were modelled as Poisson spike generators, with evoked firing determined by a receptor response profile and a spontaneous baseline. In the absence of stimulus the spontaneous firing rate of all ORNs is set to 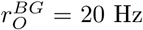. In the presence of a stimulus the ORN firing rate is given by the summation of the spontaneous rate and an activation Δ*r_O_*:

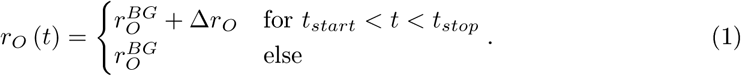

The intensity (amplitude) of ORN activation Δ*r_O_* is given by the receptor response profile that depends on receptor type and stimulus identity. Receptor activation follows a sine profile over half a period (0 … *π*):

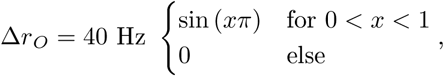

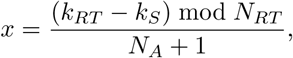

where *k_S_* is the stimulus index, *k_RT_* the receptor type index, *N_rt_* = 35 is the total number of receptor types and *N_a_* = 11 is the number of receptor types activated by a stimulus. Given these parameters 35 different odor responses can be simulated *(k_S_* = 0 … 34). This profile ensures that odor responses are evenly distributed across receptor types, while the choice of the sine shape was arbitrary. If the difference between the index of two stimuli Δ*k_s_* is small, those two stimuli are called similar, because they elicit largely overlapping responses. For Δ*k_s_* > 12 the responses do not overlap representing dissimilar stimuli.

#### 4.1.2 Neuron model

PNs, LNs, and KCs were modelled as leaky integrate-and-fire neurons with conductance-based synapses and a spike-triggered adaptation [82] current *I^A^.* The membrane potential of the i-th neuron from the PN, LN, and KC populations obeys:

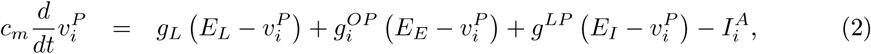

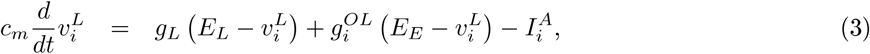

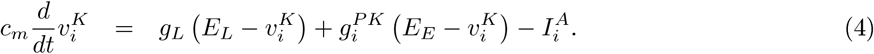

Membrane potentials follow a fire-and-reset rule. The fire-and-reset rule defines the spike trains of PNs, LNs and KCs denoted by 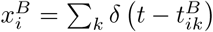 for the i-th neuron of type B. The spike trains of the ORN neurons are generated by a Poisson process with spike times 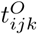 for the j-th receptor neuron of the k-th receptor type:

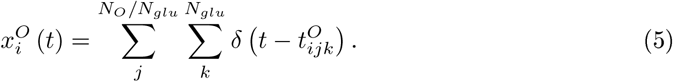

Synaptic conductances *g_i_* obey:

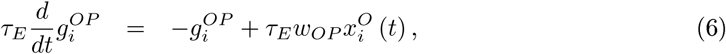

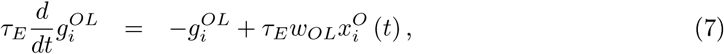

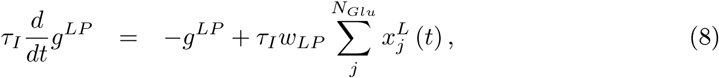

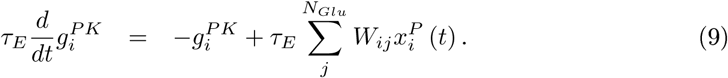

Adaptation currents 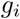 obey:

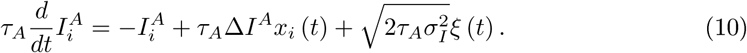

where τ_A_ is the time constant and Δ*I^A^* the spike-triggered increase of the adaptation current. The last term reflects the diffusion approximation of channel noise [40], where ξ *(t)* represents Gaussian, white noise. The variance of the adaptation currents 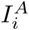 is given by 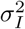.

#### 4.1.3 Simulation conditions

Four different scenarios were simulated: without lateral inhibition and cellular adaptation (i), with lateral inhibition (ii), with cellular adaptation (iii) and with lateral inhibition and cellular adaptation (iv). We quantified the strength of lateral inhibition with a multiplicative factor *α*, that set by the synaptic weight *w_LP_* in units of *w_OL_*:

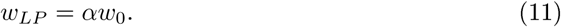

In scenarios without cellular adaptation ((i), (ii)) the dynamic adaptation current was replaced by a static current 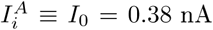 in the PN and LN populations, whereas in the KC population it was set to zero 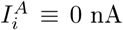. In scenarios without lateral inhibition ((i),(iii)) the inhibitory weights *w_LP_* were set to zero by setting *α* = 0.

The synaptic weight *w_OL_* was adjusted to achieve a spontaneous LN firing rate of ~ 8 Hz that is well within the experimentally observed range [1, 42].

In all scenarios the spontaneous firing rate of PNs was set to ~ 8 Hz [1, 42, 43], by adjusting the synaptic weights between the ORNs and the PNs *w_OP_*

### 4.2 Data analysis

#### 4.2.1 Population firing rate

The spike count of the i-th neuron, in the k-th time bin with size Δ*t* is given by:

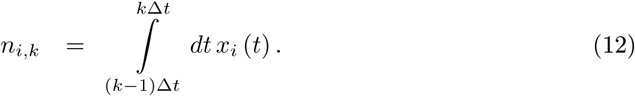

Population firing rates were obtained from the spike count in a small time bin (Δ*t* = 10 ms)

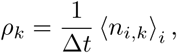

where 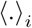 indicates the population average. In addition population firing rates were averaged over 50 trials.

#### 4.2.2 Sparseness measure

Sparseness of evoked KC responses was quantified by the widely used modified Treves-Rolls measure [47, 83]:

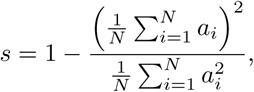

where *a_i_* indicates either the distribution of KC spike counts (population sparseness, for *i* between 1 and 1000), or binned KC population firing rate (temporal sparseness, Δ*t* = 50ms, for *i* between 1 and 20). The sparseness measure takes values between zero and one, high values indicate sparse responses. Both measures were averaged over 50 trials.

#### 4.2.3 Pattern overlap

Pattern overlap between two similar odors was calculated using Pearson’s correlation coefficient:

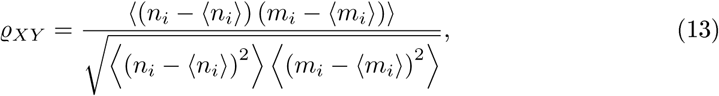

where *n_i_* and *m_i_*, are the spike count vectors of the i-th neuron in response to two respective odors (Δ*k_S_* = 2). The averages (indicated by 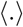) are taken over neurons. The correlation coefficient was calculated both for the PN and the KC population, and averaged over 50 trials and 5 network realizations with randomly drawn PN-KC connectivity.

##### Lateral inhibition scaling with parameter *α*

In order to test if the decrease of overlap was robust for different strengths of lateral inhibition, the synaptic weight *w_OP_* was adjusted as follows:

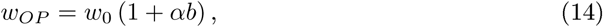

where *b* was estimated from simulations under the condition that for a range of lateral inhibition strengths (*α* ∈ [0, 9]) the spontaneous PN firing rate was close to 8 Hz.

#### 4.2.4 Decoding analysis

Odor identity was recovered from odor representations by Gaussian naive Bayes classification [84], using the scikit-learn package [85]. Training and testing data consisted of simulated odor representations for a set of seven stimuli *(k_S_* = 0, 2, …, 12), 50 trials each. Classification was repeated for every time bin (Δ*t* = 50 ms, 60 bins total) for PN spike counts, KC spike counts, or amplitudes of KC adaptations currents. Data was divided into a training and testing set using a 3-fold cross-validation procedure. Decoding accuracy was estimated by the *maximum a posteriori* method and is given by the fraction of successful classification trials divided by the total number of test trials.

### 4.3 Parameters of the neuron model

| Neuron Parameters |  |  |
| --- | --- | --- |
| membrane capacitance | $c_m$ | 289.5 pF |
| leak conductance | $g_L$ | 28.95 nS |
| leak potential | $E_L$ | -70 mV |
| reset potential | $V_R$ | -70 mV |
| threshold potential | $V_T$ | -57 mV |
| refractory time | $\tau_{ref}$ | 5 ms |
| Synaptic Parameters |  |  |
| synaptic weight | $w_0$ | 1 nS |
| excitatory synaptic potential | $E_E$ | 0 mV |
| excitatory time constant | $\tau_E$ | 2 ms |
| inhibitory synaptic potential | $E_I$ | -75 mV |
| inhibitory time constant | $\tau_I$ | 10 ms |
| Adaptation Parameters |  |  |
| spike triggered current | $\Delta I^A$ | 0.132 nA |
| adaptation time constant | $\tau_A$ | 389 ms |
| adaptation current variance | $\sigma_I^2$ | 87.1 pA<sup>2</sup> |

## Authors’ contributions

RB and MPN designed the research, RB implemented the model and analyzed the data, RB and MPN discussed results and analysis and drafted the manuscript; BL discussed results and analysis, and helped drafting the manuscript. All authors commented on the manuscript and gave final approval for publication.

## Competing interests

We declare we have no competing interests.

## Funding

RB received funding by the DFG through the Research Training Group “Sensory Computation in Neural Systems” (GRK 1589). MN received funding from them German Federal Ministry of Education and Research (BMBF) within the Bernstein Focus Neuronal Learning: “Insect Inspired Robots’ (Grant 01GQ0941).

## Acknowledgement

We thank Farzad Farkhooi for the initial network model and conceptual discussions.

## References

[1] Perez-Orive J, Mazor O, Turner GC, Cassenaer S, Wilson RI, Laurent G, 2002 Oscillations and sparsening of odor representations in the mushroom body. Science 297, 359–65. doi:10.1126/science.1070502

[2] Szyszka P, Ditzen M, Galkin A, Galizia CG, Menzel R, Ditzen M, Galkin A, Giovanni C, 2005 Sparsening and Temporal Sharpening of Olfactory Representations in the Honeybee Mushroom Bodies. J. Neurophysiol. 94, 3303–3313. doi:10.1152/jn.00397.2005.

[3] Ito I, Ong RCY, Raman B, Stopfer M, 2008 Sparse odor representation and olfactory learning. Nat. Neurosci. 11, 1177–84. doi:10.1038/nn.2192

[4] Turner GC, Bazhenov M, Laurent G, 2008 Olfactory Representations by Drosophila Mushroom Body Neurons. J. Neurophysiol. 734–746. doi:10.1152/jn.01283.2007.

[5] Honegger KS, Campbell Raa, Turner GC, 2011 Cellular-resolution population imaging reveals robust sparse coding in the Drosophila mushroom body. J. Neurosci. 31, 11772–85. doi:10.1523/JNEUROSCI.1099-11.2011

[6] Hromadka T, DeWeese MR, Zador AM, 2008 Sparse representation of sounds in the unanesthetized auditory cortex. PLoS Biol. 6, el6. doi:10.1371/journal.pbio.0060016

[7] Vinje WE, Gallant JL, 2000 Sparse coding and decorrelation in primary visual cortex during natural vision. Science 287, 1273–1276. doi:10.1126/science.287.5456.1273

[8] Wolfe J, Houweling AR, Brecht M, 2010 Sparse and powerful cortical spikes. Curr. Opin. Neurobiol. 20, 306–312. doi:10.1016/j.conb.2010.03.006

[9] Isaacson JS, 2010 Odor representations in mammalian cortical circuits. Curr. Opin. Neurobiol. 20, 328–331. doi:10.1016/j.conb.2010.02.004

[10] Laughlin SB, Sejnowski TJ, 2003 Communication in neuronal networks. Science 301, 1870–1874. doi: 10.1126/science.1089662

[11] Faisal AA, Selen LPJ, Wolpert DM, 2008 Noise in the nervous system. Nat. Rev. Neurosci. 9, 292. doi: 10.1038/nrn2258

[12] Gupta N, Stopfer M, 2012 Functional analysis of a higher olfactory center, the lateral horn. J. Neurosci. 32, 8138–48. doi:10.1523/JNEUROSCI.1066-12.2012

[13] Lin AC, Bygrave AM, de Calignon A, Lee T, Miesenbock G, 2014 Sparse, decorrelated odor coding in the mushroom body enhances learned odor discrimination. Nat. Neurosci. 17, 559–68. doi:10.1038/nn.3660

[14] Huerta R, Nowotny T, 2009 Fast and robust learning by reinforcement signals: explorations in the insect brain. Neural Comput. 21, 2123–51. doi:10.1162/neco.2009.03-08-733

[15] Wessnitzer J, Young JM, Armstrong JD, Webb B, 2012 A model of non-elemental olfactory learning in Drosophila. J. Comput. Neurosci. 32, 197–212. doi:10.1007/sl0827-011-0348-6

[16] Ardin P, Peng F, Mangan M, Lagogiannis K, Webb B, 2016 Using an Insect Mushroom Body Circuit to Encode Route Memory in Complex Natural Environments. PLoS Comput. Biol. 12. doi:10.1371/journal.pcbi.1004683

[17] Peng F, Chittka L, 2016 A Simple Computational Model of the Bee Mushroom Body Can Explain Seemingly Complex Forms of Olfactory Learning and Memory. Curr. Biol. 0, 2597–2604. doi:10.1016/j.cub.2016.10.054

[18] Mliller J, Nawrot M, Menzel R, Landgraf T, 2017 A neural network model for familiarity and context learning during honeybee foraging flights. Biol. Cybern. 1–14. doi:10.1007/s00422-017-0732-z

[19] Schmuker M, Pfeil T, Nawrot MP, 2014 A neuromorphic network for generic multivariate data classification. PNAS 111, 2081–6. doi:10.1073/pnas.1303053111

[20] Mosqueiro TS, Huerta R, 2014 Computational models to understand decision making and pattern recognition in the insect brain. Curr. Opin. insect Sci. 6, 80–85. doi:10.1016/j.cois.2014.10.005

[21] Benda J, Herz AVM, 2003 A universal model for spike-frequency adaptation. Neural Comput. 15, 2523–64. doi:10.1162/089976603322385063

[22] Farkhooi F, Muller E, Nawrot M, 2011 Adaptation reduces variability of the neuronal population code. Phys. Rev. E 83, 1–4. doi:10.1103/PhysRevE.83.050905

[23] Farkhooi F, Froese A, Muller E, Menzel R, Nawrot MP, 2013 Cellular Adaptation Facilitates Sparse and Reliable Coding in Sensory Pathways. PLoS Comput. Biol. 9, el003251. doi:10.1371/journal.pcbi.1003251

[24] Kuffler SW, 1953 Discharge Patterns and Functional Organization of Mammalian Retina. J. Neurophysiol. 16, 37–68

[25] Hartline HK, Wagner HG, Ratliff F, 1956 Inhibition in the eye of Limulus. J. Gen. Physiol. 39, 651–73. doi:10.1085/jgp.200709918

[26] Fuchs JL, Brown PB, 1984 Two-point discriminability: Relation to properties of the somatosensory system. Somatosens. Res. 2, 163–169

[27] Oswald AMM, Schiff ML, Reyes AD, 2006 Synaptic mechanisms underlying auditory processing. Curr. Opin. Neurobiol. 16, 371–376. doi:10.1016/j.conb.2006.06.015

[28] Nagel KI, Wilson RI, 2011 Biophysical mechanisms underlying olfactory receptor neuron dynamics. Nat. Neurosci. 14, 208–16. doi:10.1038/nn.2725

[29] Krofczik S, Menzel R, Nawrot MP, 2009 Rapid odor processing in the honeybee antennal lobe network. Front. Comput. Neurosci. 2, 9. doi:10.3389/neuro.l0.009.2008

[30] Wtistenberg DC, Boytcheva M, Grtinewald B, Byrne JH, Menzel R, Baxter Da, 2004 Current- and voltage-clamp recordings and computer simulations of Kenyon cells in the honeybee. J. Neurophysiol. 92, 2589–603. doi:10.1152/jn.01259.2003

[31] Demmer H, Kloppenburg P, 2009 Intrinsic membrane properties and inhibitory synaptic input of kenyon cells as mechanisms for sparse coding? J. Neurophysiol. 102, 1538–50. doi:10.1152/jn.00183.2009

[32] Wilson RI, 2013 Early olfactory processing in Drosophila: mechanisms and principles. Annu. Rev. Neurosci. 36, 217–41. doi:10.1146/annurev-neuro-062111-150533

[33] Wilson RI, Turner GC, Laurent G, 2004 Transformation of olfactory representations in the Drosophila antennal lobe. Science 303, 366–370. doi:10.1126/science.1090782

[34] Stopfer M, Jayaraman V, Laurent G, 2003 Intensity versus identity coding in an olfactory system. Neuron 39, 991–1004. doi:10.1016/j.neuron.2003.08.011

[35] Olsen SR, Wilson RI, 2008 Lateral presynaptic inhibition mediates gain control in an olfactory circuit. Nature 452, 956–960. doi:10.1038/nature06864

[36] Wilson RI, Laurent G, 2005 Role of GABAergic inhibition in shaping odor-evoked spatiotemporal patterns in the Drosophila antennal lobe. J. Neurosci. 25, 9069–79. doi:10.1523/JNEUROSCI.2070-05.2005

[37] Deisig N, Giurfa M, Sandoz JC, 2010 Antennal lobe processing increases separability of odor mixture representations in the honeybee. J. Neurophysiol. 103, 2185–2194. doi:10.1152/jn.00342.2009

[38] Capurro A, Baroni F, Olsson SB, Kuebler LS, Karout S, Hansson BS, Pearce TC, 2012 Non-linear blend coding in the moth antennal lobe emerges from random glomerular networks. Front. Neuroeng. 5, 6. doi: 10.3389/fneng.2012.00006

[39] Caron SJC, Ruta V, Abbott LF, Axel R, 2013 Random convergence of olfactory inputs in the Drosophila mushroom body. Nature 497, 113–7. doi:10.1038/naturel2063

[40] Schwalger T, Fisch K, Benda J, Lindner B, 2010 How noisy adaptation of neurons shapes interspike interval histograms and correlations. PLoS Comput. Biol. 6, el001026. doi:10.1371/journal.pcbi.1001026

[41] Fisch K, Schwalger T, Lindner B, Herz A, Benda J, 2012 Channel noise from both slow adaptation currents and fast currents is required to explain spike-response variability in a sensory neuron. J. Neurosci. 32, 17332. doi:10.1523/JNEUROSCI.6231-11.2012

[42] Chou YH, Spletter ML, Yaksi E, Leong JCS, Wilson RI, Luo L, 2010 Diversity and wiring variability of olfactory local interneurons in the Drosophila antennal lobe. Nat. Neurosci. 13, 439–49. doi:10.1038/nn.2489

[43] Meyer A, Galizia CG, Nawrot MP, 2013 Local interneurons and projection neurons in the antennal lobe from a spiking point of view. J. Neurophysiol. 110, 2465–74. doi:10.1152/jn.00260.2013

[44] Nawrot MP, 2012 Dynamics of sensory processing in the dual olfactory pathway of the honeybee. Apidologie doi:10.1007/sl3592-012-0131-3

[45] Strube-Bloss MF, Herrera-Valdez Ma, Smith BH, 2012 Ensemble response in mushroom body output neurons of the honey bee outpaces spatiotemporal odor processing two synapses earlier in the antennal lobe. PLoS One 7, e50322. doi:10.1371/journal.pone.0050322

[46] Belmabrouk H, Nowotny T, Rospars JP, Martinez D, 2011 Interaction of cellular and network mechanisms for efficient pheromone coding in moths. Proc. Natl. Acad. Sci. U. S. A. 108, 19790–5. doi: 10.1073/pnas.1112367108

[47] Treves A, Rolls ET, 1991 What determines the capacity of autoassociative memories in the brain? Netw. Comput. Neural Syst. 2, 371–397. doi:10.1088/0954-898X/2/4/004

[48] Laurent G, 2002 Olfactory network dynamics and the coding of multidimensional signals. Nature Rev. Neurosci. 3, 884

[49] Campbell Raa, Honegger KS, Qin H, Li W, Demir E, Turner GC, 2013 Imaging a population code for odor identity in the Drosophila mushroom body. J. Neurosci. 33, 10568–81. doi:10.1523/JNEUROSCI.0682-12.2013

[50] Szyszka P, Demmler C, Oemisch M, Sommer L, Biergans S, Birnbach B, Silbering AF, Galizia CG, 2011 Mind the gap: olfactory trace conditioning in honeybees. J. Neurosci. 31, 7229–39. doi:10.1523/JNEUROSCL6668-10.2011

[51] Marr BYD, 1969 A theory of cerebellar cortex. J. Physiol. 202, 437–470. doi:10.2307/1776957

[52] Albus JS, 1971 A theory of cerebellar function. Math. Biosci. 10, 25–61. doi:10.1016/0025-5564(71)90051-4

[53] Kanerva P, 1988 Sparse Distributed Memory. MIT Press, Cambridge, MA

[54] Stierle JS, Galizia CG, Szyszka P, 2013 Millisecond stimulus onset-asynchrony enhances information about components in an odor mixture. J. Neurosci. 33, 6060–9. doi:10.1523/JNEUROSCI.5838-12.2013

[55] Galili DS, Liidke A, Galizia CG, Szyszka P, Tanimoto H, 2011 Olfactory trace conditioning in Drosophila. J. Neurosci. 31, 7240–7248. doi:10.1523/JNEUROSCI.6667-10.2011

[56] Assisi C, Stopfer M, Laurent G, Bazhenov M, 2007 Adaptive regulation of sparseness by feedforward inhibition. Nat. Neurosci. 10, 1176–84. doi:10.1038/nnl947

[57] Papadopoulou M, Cassenaer S, Nowotny T, Laurent G, 2011 Normalization for sparse encoding of odors by a wide-field interneuron. Science 332, 721–5. doi:10.1126/science.1201835

[58] Lei Z, Chen K, Li H, Liu H, Guo A, 2013 The GABA system regulates the sparse coding of odors in the mushroom bodies of Drosophila. Biochem. Biophys. Res. Commun. 436, 35–40. doi:10.1016/j.bbrc.2013.05.036

[59] Kee T, Sanda P, Gupta N, Stopfer M, Bazhenov M, 2015 Feed-Forward versus Feedback Inhibition in a Basic Olfactory Circuit. PLOS Comput. Biol. 11, el004531. doi:10.1371/journal.pcbi.1004531

[60] Takahashi N, Katoh K, Watanabe H, Nakayama Y, Iwasaki M, 2017 Complete identification of four giant interneurons supplying mushroom body calyces in the cockroach Periplaneta americana. J. Comp. Neurol. 525, 204–230. doi:10.1002/cne.24108

[61] Liu X, Davis RL, 2009 The GABAergic anterior paired lateral neuron suppresses and is suppressed by olfactory learning. Nat. Neurosci. 12, 53–59. doi:10.1038/nn.2235

[62] Grtinewald B, 1999 Morphology of feedback neurons in the mushroom body of the honeybee, Apis mellifera. J. Comp. Neurol. 404, 114–126

[63] Froese A, Szyszka P, Menzel R, 2014 Effect of GABAergic inhibition on odorant concentration coding in mushroom body intrinsic neurons of the honeybee. J. Comp. Physiol. A 200, 183–195. doi:10.1007/s00359-013-0877-8

[64] Jortner RA, Farivar SS, Laurent G, 2007 A Simple Connectivity Scheme for Sparse Coding in an Olfactory System. J. Neurosci. 27, 1659–1669. doi:10.1523/JNEUROSCI.4171-06.2007

[65] Haehnel M, Menzel R, 2010 Sensory Representation and Learning-Related Plasticity in Mushroom Body Extrinsic Feedback Neurons of the Protocerebral Tract. Front. Syst. Neurosci. 4, 1–13. doi: 10.3389/fnsys.2010.00161

[66] Haenicke J, 2015 Modeling insect inspired mechanisms of neural and behavioral plasticity. Ph.D. thesis

[67] Filla I, Menzel R, 2015 Mushroom body extrinsic neurons in the honeybee (Apis mellifera) brain integrate context and cue values upon attentional stimulus selection. J. Neurophysiol. 114, 2005–2014. doi: 10.1152/jn.00776.2014

[68] Barlow H, 1961 Possible principles underlying the transformations of sensory messages. In Sens. Commun.: vol. 6, 57–58. doi:10.7551/mitpress/9780262518420.003.0013

[69] Barlow H, 2001 Redundancy reduction revisited. Netw. Comput. Neural Syst. 12, 241–253. doi: 10.1080/net.l2.3.241.253

[70] Uchida N, Poo C, Haddad R, 2013 Coding and Transformations in the Olfactory System. Annu. Rev. Neurosci. 363–385. doi:10.1146/annurev-neuro-071013-013941

[71] Friedrich RW, Wiechert MT, 2014 Neuronal circuits and computations: pattern decorrelation in the olfactory bulb. FEBS Lett. doi:10.1016/j.febslet.2014.05.055

[72] Galizia CG, 2014 Olfactory coding in the insect brain: data and conjectures. Eur. J. Neurosci. 1–12. doi: 10.1111/ejn.12558

[73] Luo SX, Axel R, Abbott LF, 2010 Generating sparse and selective third-order responses in the olfactory system of the fly. PNAS 107, 10713–8. doi:10.1073/pnas.1005635107

[74] Bhandawat V, Olsen SR, Gouwens NW, Schlief ML, Wilson RI, 2007 Sensory processing in the Drosophila antennal lobe increases reliability and separability of ensemble odor representations. Nat. Neurosci. 10, 1474–82. doi:10.1038/nnl976

[75] Olsen SR, Bhandawat V, Wilson RI, 2010 Divisive normalization in olfactory population codes. Neuron 66, 287–299. doi:10.1016/j.neuron.2010.04.009

[76] Mazor O, Laurent G, 2005 Transient dynamics versus fixed points in odor representations by locust antennal lobe projection neurons. Neuron 48, 661–73. doi:10.1016/j.neuron.2005.09.032

[77] Perisse E, Waddell S, 2011 Associative memory: Without a trace. Curr. Biol. 21, R579–R581. doi: 10.1016/j.cub.2011.06.012

[78] Dylla KV, Galili DS, Szyszka P, Ludke A, 2013 Trace conditioning in insects-keep the trace! Front. Physiol. 4 AUG, 1–12. doi:10.3389/fphys.2013.00067

[79] Buonomano DV, Maass W, 2009 State-dependent computations: spatiotemporal processing in cortical networks. Nat. Rev. Neurosci. 10, 113–125. doi:10.1038/nrn2558

[80] Liidke A, Raiser G, Nehrkorn J, Herz AVM, Galizia CG, Szyszka P, 2018 Calcium in Kenyon Cell Somata as a Substrate for an Olfactory Sensory Memory in Drosophila. Front. Cell. Neurosci. 12, 128. doi: 10.3389/fncel.2018.00128

[81] Goodman DFM, Brette R, 2009 The Brian Simulator. Front. Neurosci. 3, 192–197. doi: 10.3389/neuro.01.026.2009

[82] Treves A, 1993 Mean-field analysis of neuronal spike dynamics. Network: Comput. Neural Syst. 4, 259. doi:10.1088/0954-898X/4/3/002

[83] Willmore B, Tolhurst DJ, 2001 Characterizing the sparseness of neural codes. Network 12, 255–270. doi: 10.1088/0954-898X/12/3/302

[84] Rish I, 2001 An empirical study of the naive Bayes classifier. In Work. Empir. methods Artif. Intell., vol. 22230, 41–46

[85] Pedregosa F, et al., 2012 Scikit-learn: Machine Learning in Python. J. Mach. Learn. Res. 12, 2825–2830. doi:10.1007/sl3398-014-0173-7.2

